# dcHiC: differential compartment analysis of Hi-C datasets

**DOI:** 10.1101/2021.02.02.429297

**Authors:** Abhijit Chakraborty, Jeffrey Wang, Ferhat Ay

## Abstract

The compartmental organization of chromatin and its changes play important roles in distinct biological processes carried out by mammalian genomes. However, differential compartment analyses have been mostly limited to pairwise comparisons and with a main focus on only the compartment flips (e.g., A-to-B). Here, we introduce *dcHiC*, which utilizes quantile normalized compartment scores and a multivariate distance measure to identify significant changes in compartmentalization among multiple contact maps. Evaluating dcHiC on three collections of Hi-C contact maps from mouse neural differentiation (n=3), mouse hematopoiesis (n=10) and human LCL cell lines (n=20), we show its effectiveness and sensitivity in detecting biologically relevant differences, including those validated by orthogonal experiments. Across these experiments, *dcHiC* reported regions with dynamically regulated genes associated with cell identity, along with correlated changes in chromatin states, replication timing and lamin B1 association. With its efficient implementation, dcHiC not only enables high-resolution compartment analysis but also includes a suite of additional features, including standalone browser visualization, differential interaction identification, and time-series clustering. As such, it is an essential addition to the Hi-C analysis toolbox for the ever-growing number of contact maps being generated. *dcHiC* is freely available at https://github.com/ay-lab/dcHiC, and examples from this paper can be seen at https://ay-lab.github.io/dcHiC.

## BACKGROUND

The three-dimensional organization of chromatin in the nucleus has been of interest to scientists for more than a century now. The observation that different chromosomes occupy a defined space in the nucleus dates back to Carl Rabl’s work in animal cells in 1885 [1]. Since then, many experimental techniques have been developed to image and map chromatin, allowing us to look at chromatin organization at an ever-increasing resolution. The greatest strides in this area have been made in the past decade following the advent of genome-wide conformation capture techniques. We now know that interphase chromosomes are folded into multiple layers of hierarchical structures. Each layer contributes to the establishment and maintenance of the epigenetic landscape that controls cellular state and function.

Among these, the megabase-scale compartmental organization of eukaryotic genomes has been shown to play a critical role in transcription, DNA replication, accumulation of mutations, and DNA methylation [2-12]. In broad terms, two types of compartments divide the genome into regions of open and active chromatin (compartment A) versus inactive and closed chromatin (compartment B) [13]. Further analysis of each compartment revealed subsets of regions with markedly different properties within each class called subcompartments [14, 15] as well as to a putative third class (intermediate or I) that is at the interface between A and B and is reorganized in tumors [9].

The main method to extract compartment information has been to analyze high-throughput chromosome conformation capture (Hi-C) contact maps using Principal Components Analysis (PCA) [13, 16, 17]. Briefly, this process involves distance normalization (observed/expected for each genomic distance) of the Hi-C contact map for each chromosome at a particular resolution (generally between 100 kb to 1 Mb) followed by transformation into a correlation matrix, where each entry (*i, j)* denotes the correlation of row *i* and row *j* (or column *i* and *j* since symmetric) of the distance-normalized Hi-C map. The eigenvalue decomposition of the correlation matrix provides the eigenvectors, and the first eigenvector or principal component (PC1) typically represents the genomic compartments A and B. If PC1 corresponds to chromosome arms or other broad patterns in the Hi-C map (e.g., copy number differences), the second principal component (PC2) is likely to represent A and B compartments. The A/B compartment labels are assigned to the positive/negative stretches of the selected PC; however, depending on the implementation of eigenvalue decomposition, it may be necessary to reorient these assignments correctly using GC content or gene density.

Whether one is interested in the two major compartments or their more nuanced subsets, the magnitude and sign of eigenvalues derived from PCA have been the major determinants of compartment type. However, standard PCA is limited in analyzing each Hi-C contact map individually, and to date, there is no method to systematically compare compartmentalization across multiple (>2) Hi-C datasets. This is becoming an obstacle in analyzing the ever-increasing chromatin conformation data, either from Hi-C or its variants [18-25], generated across many cell types and conditions [26]. Technical challenges such as selecting the correct PC and sign that represents A/B compartments and their scaling across different datasets become larger problems when comparing many Hi-C contact maps. Thus far, comparative compartment analysis has been mainly limited to examining compartment flips between two Hi-C maps at a time [27, 28].

Here, we introduce *dcHiC* (<u>d</u>ifferential <u>c</u>ompartment analysis of <u>Hi</u>-<u>C</u>), a method that identifies statistically significant differences in compartmentalization among two or more contact maps, including changes that are not accompanied by a compartment flip. Our method implements a memory-efficient and parallelized singular value decomposition (SVD) to derive principal components (i.e., eigenvectors) followed by quantile normalization to obtain comparable compartment scores across two or more Hi-C maps at a time **(Figure 1, Step 1)**. *dcHiC* then utilizes the normalized component scores to derive a multivariate distance measure [29] **(Figure 1, Step 2)** to estimate the statistical significance of compartment differences. If available, dcHiC utilizes variance among Hi-C replicates as covariates for Independent Hypothesis Weighting (IHW) [30] to correct for multiple testing. With our methodology, compartment analysis can be conducted on Hi-C maps with or without replicates at resolutions up to 10 kb for human and mouse genomes. Further downstream, dcHiC provides a raft of analysis features, including standalone IGV browser [31] visualization of results, detection of differential interactions involving significant differential compartments, time-series clustering of compartment scores, and a module for determining enriched Gene Ontology terms from differential compartments.

**Figure 1:**
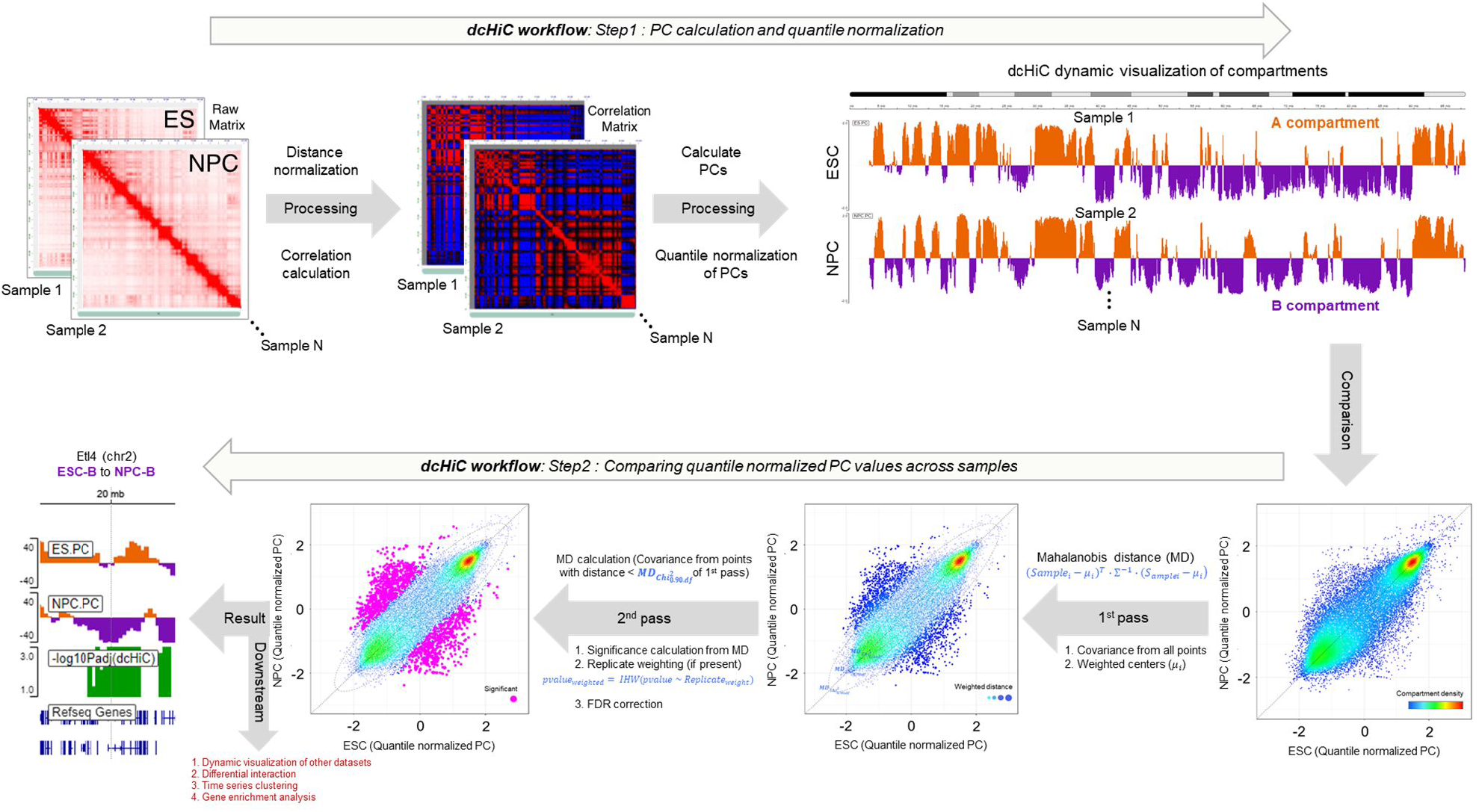
Outline of the method. The figure panel shows the *dcHiC* workflow in two steps. In step 1, *dcHiC* calculates the principal components followed by the quantile normalization of the compartment scores across all input Hi-C data. In step 2, for each genomic bin, *dcHiC* calculates a Mahalanobis distance, which is a statistical measure of the extent to which each bin is a multivariate outlier with respect to the overall multivariate compartment score distribution across all input Hi-C maps **(Methods)**. dcHiC then utilizes the Mahalanobis distance to assign a statistical significance using the chi-square test (p-value) for each compartment bin and employs independent hypothesis weighting (IHW – when there are replicate samples) or FDR (when no replicates are available) correction on these p-values. *dcHiC* outputs a standalone dynamic IGV web browser view and enables the user to integrate other datasets into the same view for an integrated visualization.

To assess the biological relevance of the identified differences, we applied *dcHiC* to several different collections of Hi-C datasets across various biological conditions, including mouse neuronal development (n=3), mouse hematopoiesis (n=10), and a set of lymphoblastoid cell lines (LCLs) from different human populations (n=20). Analyzing each Hi-C dataset at 100 kb and 40 kb resolutions, we identified relevant compartmentalization differences reflecting the underlying biology in the respective scenarios. In the mouse neuronal differentiation model, *dcHiC* identified compartmental changes for loci involving critical genes associated with cellular identities in mouse embryonic stem cells (mESC) and neuronal differentiation, such as *Dppa2/4, Zfp42, Ephb1*, and *Ptn*, as well as GO term enrichments consistent with these cellular identities. In a ten-way comparison (n=10) of key cell types from mouse hematopoiesis, across stem cells, progenitor cells, and terminally differentiated cells, *dcHiC* revealed significant compartmental changes involving key genes such as *Sox6, Meis1, Runx2, Klf5*, and many others. Across both neural and hematopoietic differentiation models, our results also highlight the importance of generally ignored compartmentalization changes within the same compartment type (within A or within B - **Figure 1**). We also demonstrate the biological significance of our differential calls through strong correlations with cell-type specific differences in lamin B1 association, histone modifications and gene expression. For human LCLs, comparing twenty Hi-C maps from a diverse set of donors, *dcHiC* confirmed the previous findings, with significant enrichment of various biological signals within the differential compartments across the population.

Overall, *dcHiC* provides an integrative framework and an easy-to-use tool for comparative analysis of Hi-C maps and identifies biologically relevant differences in compartmentalization across multiple cell types. With immediate application to hundreds of publicly available Hi-C datasets, *dcHiC* will play an essential role in providing deeper insights into dynamic genome organization and its downstream effects.

## RESULTS

### dcHiC identifies compartments consistent with the PCA-based approach

As more complex experimental designs emerge that compare different Hi-C profiles, a comprehensive method to compare the spatial organization of the genome is necessary. To do this, *dcHiC* first employs a time- and memory-efficient R implementation of singular value decomposition (SVD) to achieve the eigenvalue decomposition of each Hi-C contact map [32]. This is followed by automated selection to find the principal component and its sign (reoriented if needed) that best correlates with gene density and GC content per sample **(Methods)**. The resulting compartment scores are quantile normalized, and a multivariate score (Mahalanobis distance) is computed based on an initial covariance estimation. We then refine the null distribution by removing outliers before calculating new covariance estimates that will be used for computing the final statistical significance (Chi-square test) of differences in compartmentalization **(Methods)**. dcHiC provides standalone browser visualization as well as several other features facilitating the interpretation of its results. **Figure 1** summarizes the overall workflow of *dcHiC*.

To establish the validity of the *dcHiC* results, we first compared our implementation of the eigenvalue decomposition to commonly used PCA-based approaches, a representative of which is implemented in HOMER[33]. Beyond a few differences in prefiltering of low coverage regions, the resulting compartment scores were highly similar between *dcHiC* and HOMER for the 100 kb resolution (replicates combined) mouse ESC (Pearson’s *r=0*.*98*, **Figure 2A)** and for the mouse neuronal progenitor cell (NPC) Hi-C map (Pearson’s *r=0*.*96*, **Figure 2D**). Similar to A/B compartment decomposition from Hi-C data, association with the nuclear lamina (or radial position) is another strong indicator of a broad-level chromatin state with heterochromatin localizing at the periphery and euchromatin at the nucleus center. Such organization is a conserved feature of eukaryotic genomes across most cell types except special cases [34, 35]. Here, we used lamin B1 association profiles of ESC and NPC cell types as an independent measure of compartmentalization and compared the lamin B1 signal distribution with dcHiC and HOMER scores. As expected, both our compartment scores and HOMER results showed a strong negative correlation with lamin B1 association, confirming the previous findings [27, 36] **(Figure 2B-C, 2E-F)**. We further plotted the chromosome 16 compartment score of ESCs and NPCs from the *dcHiC*, HOMER, and lamin B1 association signal. **Figure 2G-H** shows the lamin B1 signal and compartment features captured by dcHiC and HOMER at a genome-wide scale in ESC and NPC cell types. These results establish that dcHiC, similar to the existing PCA-based (HOMER) approach, accurately captures compartment patterns.

**Figure 2:**
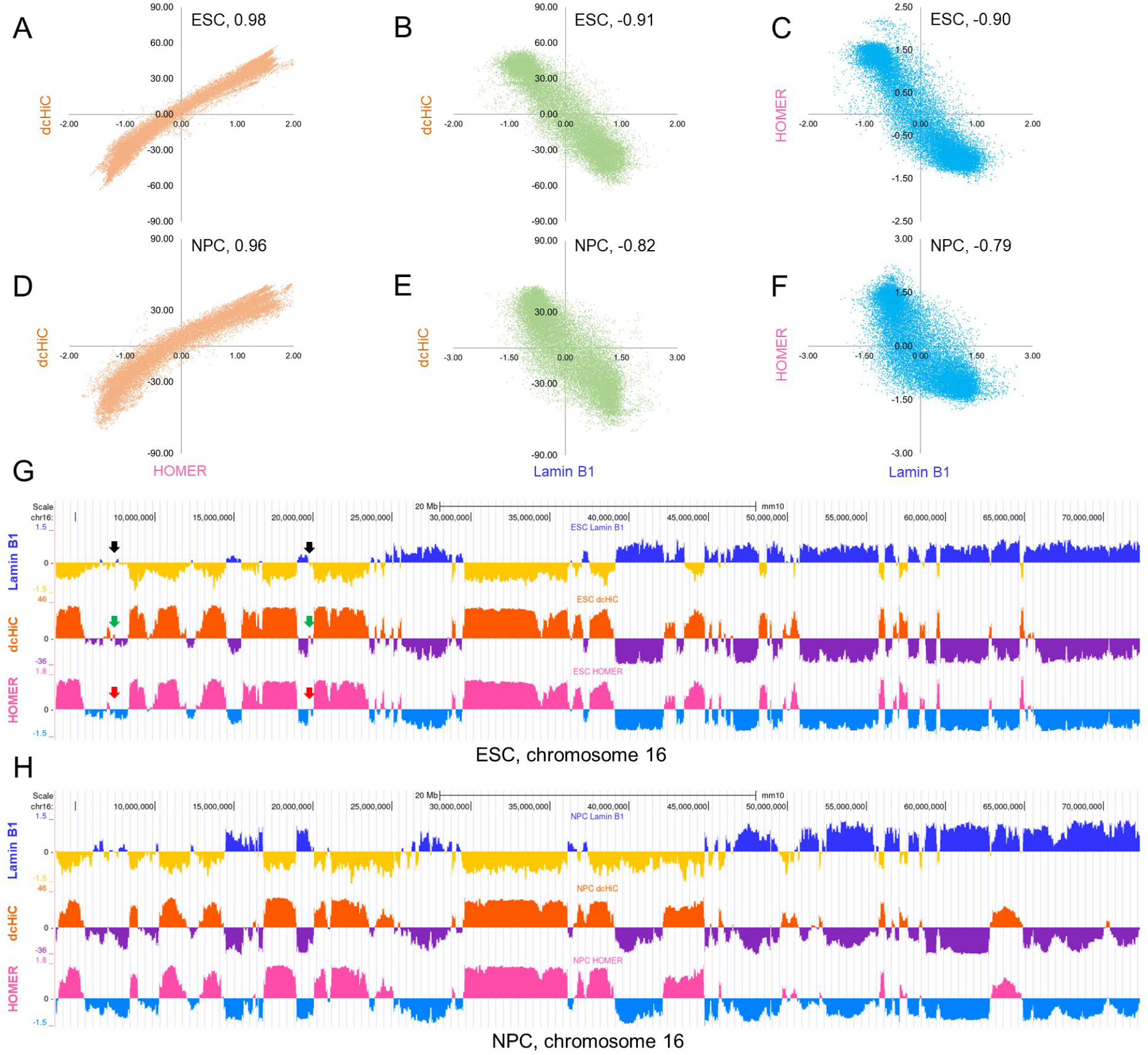
Comparison of *dcHiC* compartment scores with HOMER compartment scores and Lamin B1 association data. **(A-B)** Genome-wide comparison of *dcHiC* compartment scores against HOMER compartment scores and Lamin B1 profiles for mouse ESCs. **(C)** Comparison between HOMER compartment scores and Lamin B1 association. **(D-F)** Plots similar to **(A-C)** but for NPC Hi-C data. The Pearson correlation values between the two axes are reported for each plot. **(G-H)** Browser views of the compartment scores and Lamin B1 signal for a chromosome 16 region with arrows pointing to some small differences among the compartment scores of *dcHiC* and HOMER.

### Pairwise differential compartment analysis of the mouse neuronal differentiation model

Previous studies have reported substantial compartment flips during the mouse embryonic cell (ESC) to neuronal progenitor cell (NPC) transition, a well-studied *in vitro* differentiation system [37, 38]. These differences have been studied further using replication timing profiling, lamin B1 association mapping, and fluorescence *in situ* hybridization (DNA FISH) [8, 27, 36]. Therefore, we chose these two cell types to demonstrate *dcHiC’s* utility in a pairwise comparison to replicate known compartment flips and identify significant changes that do not involve flips from one compartment type to another. We also compared differential compartment calls from *dcHiC* and HOMER in this pairwise setting since HOMER does not readily allow multiway comparisons.

Overall, *dcHiC* identified 1981 100 kb bins with statistically significant differential compartmentalization (FDR < 0.1), covering up to 7.5% of the genome. For ESC and NPC, these differences constituted ∼37% (72.8 Mb) and ∼51% (101.6 Mb) of A (active) compartments, respectively. The differential compartments are further subdivided into flipping (A→B or B→A) or matching (A→A or B→B) compartment transitions. We observed that ∼74% of all the differential compartments were flips from A to B (∼30%) or B to A (∼44%) compartments during ESC to NPC transition, whereas the remaining ∼26% were within matching compartments **(Figure 3A)**. We further classified significant changes within the same compartments (A to A or B to B) based on whether the compartment scores were higher in ESC or NPC **(Figure 3B-E)**. For the resulting set of six different types of differential compartments, we plotted the distributions of compartment scores **(Figure 3B)**, lamin B1 association **(Figure 3C)**, replication timing **(Figure 3D)** and gene expression **(Figure 3E)**. As expected, more euchromatic compartments were associated with lower lamin B1 attachment, early replication timing and higher gene expression. These trends were consistent for compartment flips as well as changes within matched compartments (e.g., strong A in ESCs to weak A in NPCs).

**Figure 3:**
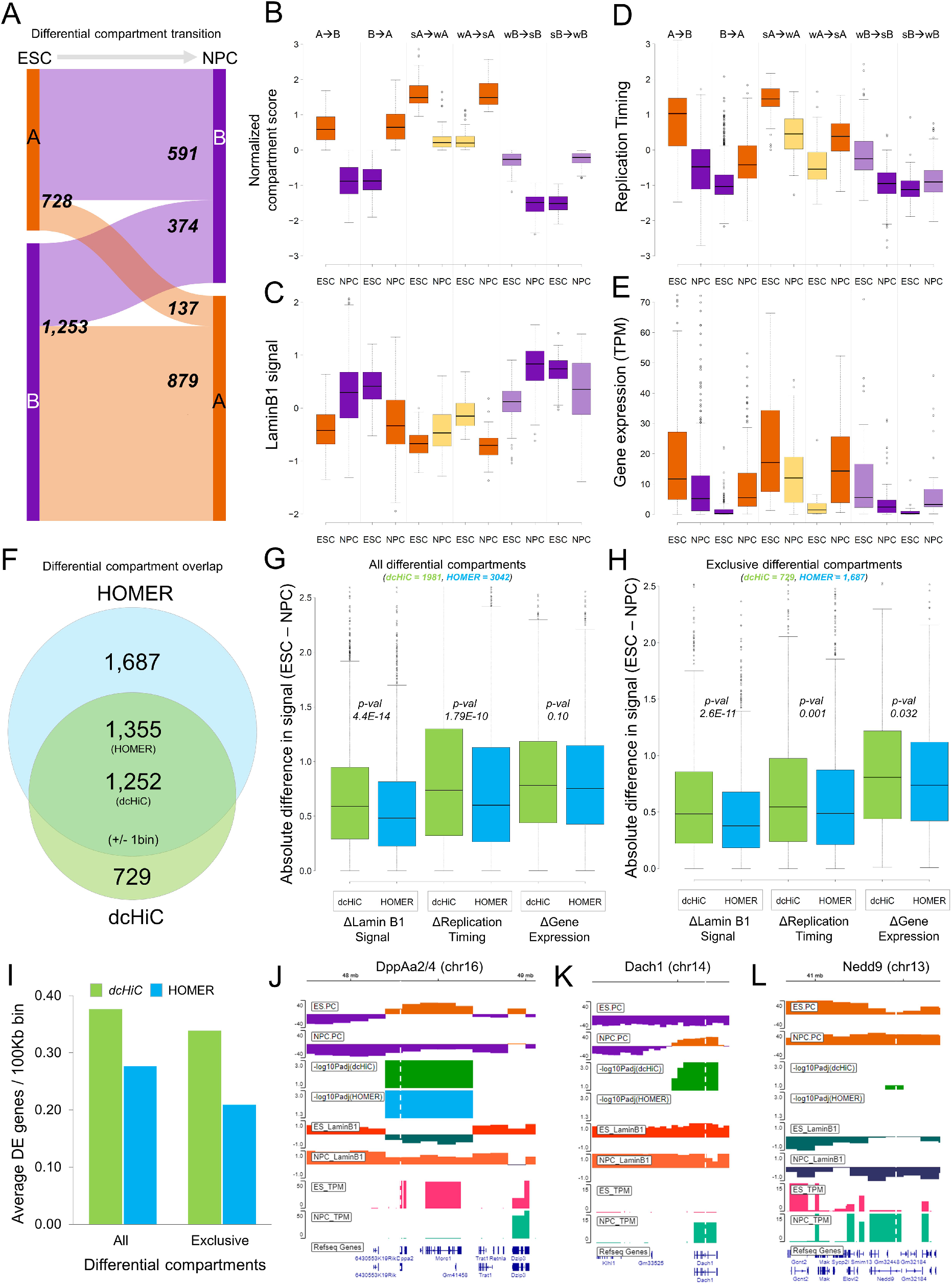
Pairwise differential compartment analysis between ESCs and NPCs. **(A)** The breakdown of the numbers of differential compartment calls (100 kb resolution) belonging to different types. **(B-E)** The distributions of compartment scores, Lamin B1, replication timing, association and gene expression across different subtypes of differential compartment calls made by *dcHiC*. Strong (s) and weak (w) were used to indicate the relative compartment strength (or absolute value) between the two cell types. **(F)** The Venn diagram shows the overlap between *dcHiC* and HOMER differential compartment calls. **(G-H)** The absolute difference in Lamin B1, replication timing signal and gene expression values (TPM) overlapping with all and exclusive differential compartments identified by *dcHiC* and HOMER, respectively (statistical significance was calculated by unpaired T-test). **(I)** The average number of differentially expressed (DE) genes overlapping with differential compartment bins (100 kb) identified by *dcHiC* and HOMER. **(J-L)** *dcHiC* differential compartments involving three DE genes: *DppA2/4, Dach1* and *Nedd9*.

Next, we compared the differential ESC vs NPC compartments from dcHiC to those from HOMER. HOMER reported a total of 3,042 100 kb bins with significant differential compartmentalization (FDR < 0.05). Only 1,355 of these 100 kb bins overlapped with dcHiC differential calls (+/- 1 bin slack; **Figure 3F**). To compare the calls made by the two different methods, we plotted the absolute differences in laminB1 signal, replication timing and log2 gene expression values of the reported differential compartments. **Figure 3G** shows the absolute difference distribution of all the respective signals from all the differential compartments between ESCs and NPCs, while **Figure 3H** shows the same but only for differential compartments exclusively identified by one method. These results show that dcHiC differential compartments are significantly (unpaired t-test p-values < 0.05) enriched for regions with higher ESC and NPC differentials for lamin association and replication timing signals. We also performed differential expression analysis between ESCs and NPCs to map the differentially expressed (DE) genes (DEseq2[39], FDR<0.05, fold change>4) on the differential compartments. We observed that *dcHiC* differential compartment bins were enriched in DE genes compared to HOMER (**Figure 3I**). The trend was similar for bins reported exclusively by each method (**Figure 3I**). These observations imply that the differential calls made by dcHiC are accompanied by larger changes between ESCs and NPCs in other biological signals relevant to compartmentalization.

To also show the utility of our tool in detecting differences at higher resolution, we ran *dcHiC* at 10 kb resolution to call differential compartments between ESC and NPC. We found a total of 16,581 10 kb bins, i.e., 165.81 Mb differential compartments between the conditions. Among the 1,981 100 kb *dcHiC* differential bins, 72% exactly overlapped at least one 10 kb differential bin (over 86% within +/-200 kb). This suggests a significant overlap across resolutions but also highlights the prevalence of regions that are detectable only at higher or lower resolution compartment analysis **(Supplementary Figure 1)**. We also evaluated the potential of false positive discoveries from *dcHiC* by running it to compare replicates of the same conditions/sample. We used all four biological Hi-C replicates available for ESC in different combinations (all 1 vs 3 and 2 vs 2 combinations of splitting the replicates). When we ran *dcHiC* on these combinations, the number of significant compartment changes (i.e., false positives) ranged from 1 to 32 with a median value of 2 bins (compared to 1,981 100 kb bins when ESC was compared to NPC), suggesting a low false positive rate for identifying differential compartments. When we ran the same analysis using 10 kb bins, we identified a median value of 751 differential bins (∼0.2% of the genome), suggesting that higher resolution differential analysis may be more prone to false positives.

#### Example genes from ESC vs NPC differential compartments

Within *dcHiC’s* calls, we also analyzed a set of key genes known for their critical role in ESC or NPC state that have been studied extensively for changes in their nuclear organization during the transition. For instance, we analyzed a set of genes for which fluorescence *in situ* hybridization (FISH) experiments were performed to study changes in radial positioning during the ESC to NPC transition. These included pluripotency markers specifically expressed in ESCs (e.g., *Zfp42* or REX1 and *Dppa2/4*) as well as EPH Receptor B1 (*Ephb1*) and other marker genes specific to neuronal differentiation. **Figure 3J** shows the *Dppa2/4* region in mouse chromosome 16 that is shown to change radial positioning, chromatin state, lamin B1 association and replication timing during differentiation [36, 40]. Consistent with these data, both dcHiC and HOMER reported a significant shift from the A (active) to the B (inactive) compartment during mouse ESC to NPC differentiation **(Figure 3J)**. In addition, *dcHiC* reported significant compartment changes for several other important genes that HOMER missed. **Figure 3K-L** displays two genes, namely, *Dach1* and *Nedd9*, which are known to play a critical role in organogenesis and signal transduction pathways for mouse neuronal development [41, 42]. We also detected these genes in our differential gene expression analysis of ESCs vs NPCs as significantly upregulated in NPCs (FDR<0.05; *>160x for Dach1* and >30x for *Nedd9*). *Dach1* lies in a compartment reported to be flipped from ESC-B to NPC-A by *dcHiC* (**Figure 3K**). *Nedd9* gene overlapped with the A compartment in both cell types but with stronger compartmentalization in NPC, which was detected as a significant change by *dcHiC* **(Figure 3L)**.

To determine whether the compartmental changes are accompanied by specific differences in local chromatin interactions, we implemented an extension of our comparative approach to identify differences in contact counts involving the differential compartments **(Methods)**. This feature allows users to input a set of significant chromatin interactions (e.g., from Fit-Hi-C[43]) or chromatin loops (e.g., from HiCCUPS or Mustache), which will then be filtered for their overlap with differential compartments and tested for their difference across the compared conditions. The black square boxes in **Figure 4A** represent the *dcHiC*-identified differential interactions (ESC vs NPC) that are anchored in the *DppA2/4* region. These interactions are identified among FitHiC2 calls [44] (FDR < 0.05) that are reported as significant in at least one replicate of ESC and/or NPC datasets. The results show that the *Dppa2/4* domain in NPC specifically interacts with its upstream region compared to ESC, while the interactions with the adjacent downstream region remained unchanged, a change that can be visualized on the Hi-C map **(Figure 4A)**. Previous studies on *Ephb1* have demonstrated significant subnuclear repositioning of the gene from the periphery to the nuclear center during ESC to NPC differentiation [36] accompanied by higher gene expression later. A similar analysis of the *Ephb1* region shows that it has enriched interactions with a pair of upstream B compartments in ESCs, which are weakened in NPCs where *Ephb1* is transitioned to the A compartment **(Figure 4B)**. In addition, the same region gained interactions with a downstream A compartment in NPC. These results highlight the value of differential interaction analysis coupled with differential compartmentalization to better delineate important changes in the local chromatin environment. Finally, even though the above examples highlight cases where gene expression is tightly correlated with compartment changes and radial positioning, this is not necessarily the case for all genes. **Figure 4C** shows the pluripotency marker gene *Pou5f1/Oct4* region with ESC-specific gene expression. The radial positioning of this gene locus was shown to remain unchanged during the ESC to NPC transition[36], consistent with our results **(Figure 4C)**. Overall, dcHiC identified both known compartment flips (A to B or B to A) as well as novel compartmentalization differences within the same compartment for important genes.

**Figure 4:**
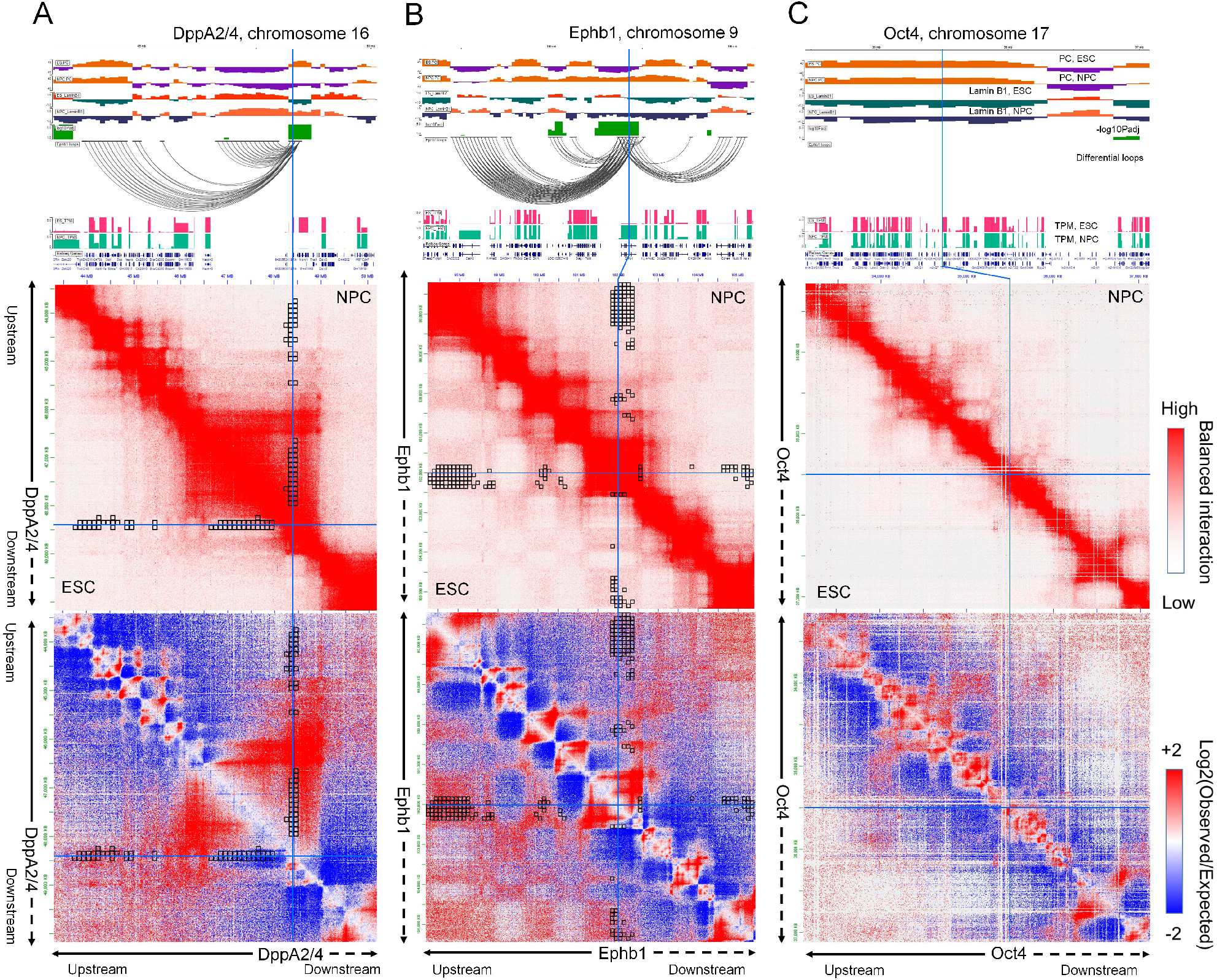
Differences in local chromatin interactions of differential compartments. Detailed browser views (top), Hi-C contact maps (mid) and differential chromatin interactions (bottom) of three gene loci – **(A)** *DppA2/4*, **(B)** *Epbh1* and **(C)** *Oct4*. Visible changes in interactions involving the *Dppa2/4* locus and *Ephb1* locus are highlighted through each plot. Although *Oct4* shows a dramatic change in gene expression, the region does not alter its radial position within the nucleus (FISH experiments), which is also consistent with the lack of change in compartmentalization reported by *dcHiC*.

### Multicell-type differential compartment analysis of the mouse neuronal system

The same *in vitro* system used to differentiate from ESCs to NPCs also allows further differentiation of NPCs to cortical neurons or CNs [38]. This developmental lineage provides an approach to demonstrate how *dcHiC* uses a multivariate distance measure to compare the compartmentalization of more than two cell types simultaneously. For such multiway comparisons, *dcHiC* provides a quick and straightforward approach to detect outliers in compartment scores and associated differential interactions, an approach far easier with many experiments than the traditional paradigm of taking pairwise comparisons. In this section, we first illustrate the biological significance of *dcHiC’s* differential compartments using multiple lines of biological data. We then demonstrate functional term enrichments and show specific differential genes that illustrate the application’s breadth of analysis.

Applying *dcHiC* at 100 kb resolution to intrachromosomal Hi-C data from ESC, NPC, and CN samples, we identified a total of 5,055 significant differential bins covering approximately 19.2% of the genome. Compartments A and B were evenly split for NPC and CN, whereas ESC had ∼63% B compartments. Overall, regions in the B compartment for each cell type were more likely to exhibit statistically significant compartment changes compared to the A compartment (21-23% vs 16-18%). **Figure 5A** summarizes the number of differential compartment bins that involve flips (A → B or B → A) or remained within the same compartment throughput the lineage transition. Consistent with the literature [2, 5, 45], we showed that compartmental dynamics are strongly associated with the variability of gene expression and histone modifications **(Figure 5B, Methods)**.

**Figure 5:**
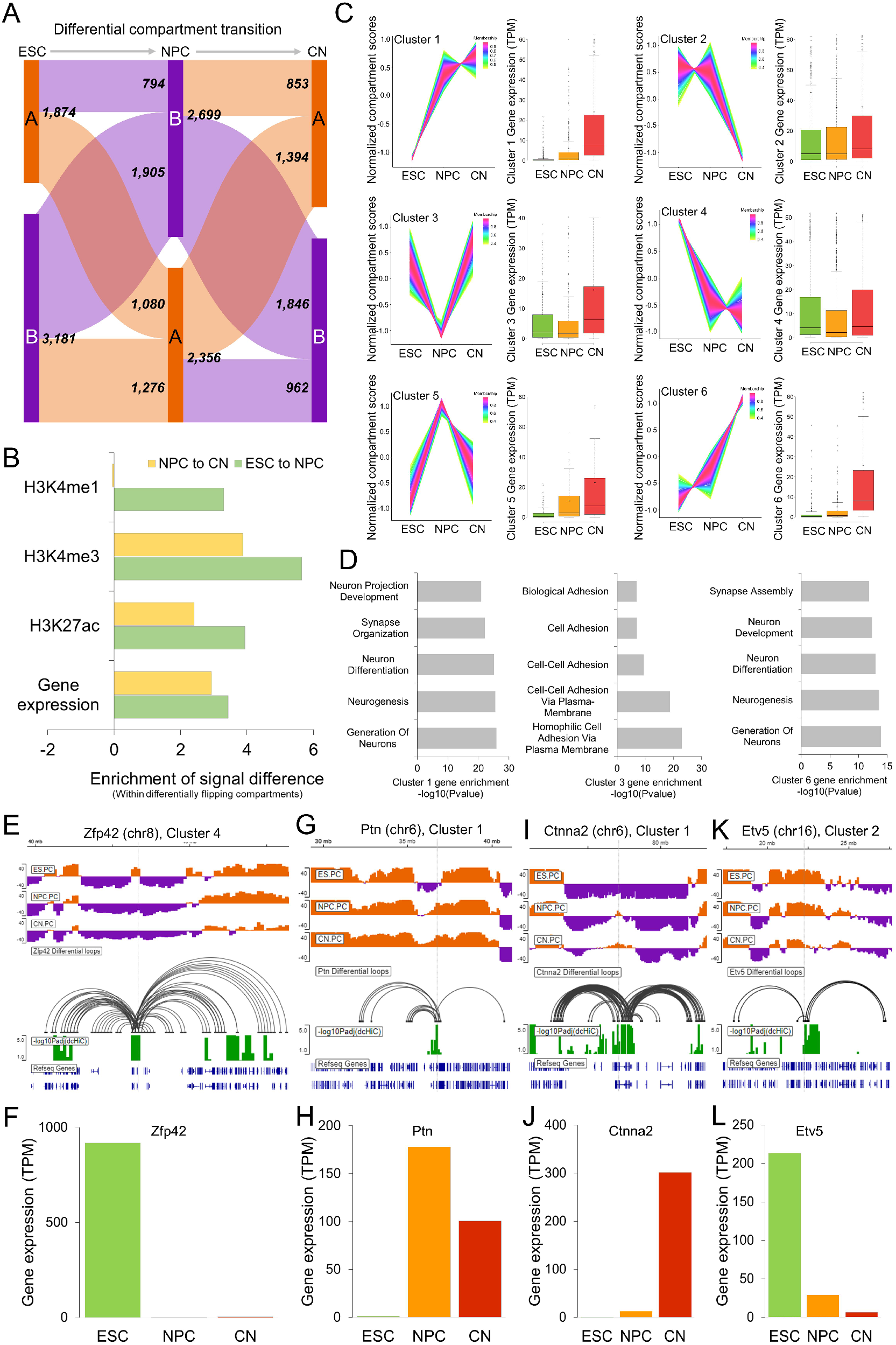
Three-way differential compartment analysis of ESCs, NPCs, and CNs. **(A)** The breakdown of the numbers of differential compartment calls belonging to different types. **(B)** The enrichment of signal differences in different histone marks and gene expression in *dcHiC* differential bins with compartment flips (A → B or B → A) compared to bins with nonsignificant compartment flips. **(C)** Time-series clustering of normalized compartment scores into six different clusters from the three cell types along with their overlapping gene expression profile. For the clustering analysis, the quantile normalized PCA scores for each 100 kb bin across ESC-NPC-CN were further z-transformed to focus on relative changes in compartmentalization. **(D)** Gene term enrichment results of GO biological functions from genes overlapping with clusters 1, 3 and 6 compartments. **(E-L)** Differential compartments overlapping with representative genes in each of the cell types shown along with the differential chromatin interactions involving the respective compartments and gene expression values (TPM) across the ESC-NPC-CN transition for four example genes, each representing one cluster pattern.

To further analyze these changes simultaneously, rather than one transition (or pair) at a time, we utilized time-series analysis to cluster the compartmentalization score patterns of these differential bins across **(Figure 5C)** and plotted the expression pattern of the overlapping genes in each cluster across three different time-points. To focus on relative changes in compartmentalization, we further z-transformed the quantile normalized PCA scores for each 100 kb bin across the three cell types and applied TC-seq[46] to identify 6 major clusters **(Methods)**. Two major clusters corresponded to regions that progressively became more euchromatic (clusters 1 and 6), and one corresponded to more heterochromatic regions (cluster 4). We observed other clusters that corresponded to one cell type showing highly different compartmentalization with respect to the other two (e.g., clusters 3 and 5 with NPC-specific patterns). To link these compartmentalization patterns to gene function, we identified genes overlapping with each differential compartment bin for each cluster. Performing functional enrichment analysis on these gene sets[47], we identified signatures that are consistent with the cellular identity of the cell type with the highest compartment z-scores (i.e., more euchromatic). For instance, for the genes overlapping with clusters 1 and 6 with compartment scores increasing from ESC to NPC to CN, the enriched terms included neurogenesis and neuronal development **(Figure 5D)**. For cluster 3, where CN compartment scores were highest, the enriched terms (cell-cell adhesion, biological adhesion, and others) were consistent with a general pattern for genes involved in regulating cell-type specific migration and development. We also observed that cluster 3 overlapped with an important class of gene family known as protocadherins [48]. Protocadherins are highly conserved genes across species, and most of them are clustered in a single genomic locus in vertebrates [49]. They are shown to be differentially expressed in individual neurons and involved in diverse neurodevelopmental processes [50]. When we repeated the functional enrichment analysis per cell type using genes overlapping A compartments with the highest compartmentalization score for that cell type compared to the other two, we also observed cellular identity-related annotation terms **(Supplementary Data 1)**. While annotations related to cell adhesion were enriched in ESCs as well as CN, CN specifically showed enrichment for neurogenesis, neuron differentiation and development **(Supplementary Data 1)**. CN, but not NPC, also showed enrichment for synaptic signaling, synapse organization and neuron projection development, potentially related to its further differentiated state with respect to NPC.

#### Example genes from ESC-NPC-CN differential compartments

The differential compartments captured by *dcHiC* encompass a variety of traditionally studied as well as more nuanced scenarios. For instance, similar to *Dppa2/4, Zfp42/Rex1* is a well-studied pluripotency marker primarily expressed in undifferentiated stem cells **(Figure 5E)**. As is the case for *Dppa2/4, Zfp42* is also in a small A compartment region surrounded by large stretches of B compartments in ESCs. As expected, this region flipped into the B compartment in NPC and stayed in CN, consistent with the lack of gene expression in these two cell types **(Figure 5F)**. *Ptn* or pleiotrophin, on the other hand, exhibits mitogenic and trophic effects on dopaminergic neurons and is instead a marker gene for neuronal lineage. *dcHiC* reported this gene in a differential compartment that is B in ESC but A in NPC and CN, in concordance with gene expression **(Figure 5G-H)**, which fits the compartmentalization pattern of cluster 1 **(Figure 5C)**. These two examples represent strong compartment flips from A to B or B to A. An example of a more gradual compartmental change is the CN-specific *Ctnna2* gene, which functions as a linker between cadherin adhesion receptors and the cytoskeleton to regulate cell-cell adhesion and differentiation in the nervous system. The B compartment encompassing *Ctnna2* in ESCs gradually weakens during the ESC-NPC-CN transition, leading to a transcription-permissive A compartment that starts in NPCs and expands further in CNs **(Figure 5I-J)**.

Compartment shifts within the same compartment are also captured by *dcHiC* **(Figure 5K-L)**. *Etv5* encodes a transcription factor that plays an important role in the segregation between epiblast and primitive endoderm specification during ESC differentiation [51]. *Etv5* is highly expressed in ESCs but gradually loses its expression **(Figure 5L)** as well as strong compartmentalization during the ESC-NPC-CN transition while remaining in the A compartment at all times. This locus belongs to cluster 2 with enrichment for more euchromatic association specifically in ESCs, consistent with the highest expression for *Etv5* for this cell type. Beyond *Etv5*, we also found a list of 199 other genes within the A compartment throughout the ESC-NPC-CN transition, for which the variation in the expression profile strongly correlated with changes in compartmentalization (Pearson correlation > 0.7; **Supplementary Data 2**). A similar analysis within differential B compartments revealed 245 genes with a strong positive correlation between expression and compartmentalization change (**Supplementary Data 2**). Overall, our results demonstrate that *dcHiC* can comprehensively analyze multiple different Hi-C maps simultaneously and identify compartmental changes involving abrupt (e.g., compartment flips) as well as gradual changes.

### Differential compartment analysis of the mouse hematopoietic system

The hematopoietic system is a developmentally regulated and well-characterized cell differentiation model [52, 53]. This system provides an opportunity to understand the dynamic changes in chromatin structure together with transcriptional and other epigenetic changes during differentiation in detail. The study of genome organization changes during this complex process—involving many different progenitors and differentiated cell types— requires a systematic approach. A recent study by Zhang et al. [28] profiled chromatin organization in a classic hematopoietic model with ten primary stem, progenitor, and terminally differentiated cell populations from mouse bone marrow **(Figure 6A)**. In this model, long-term hematopoietic stem cells (LT-HSCs) represent the starting point of the hematopoietic hierarchy with self-renewal and multilineage differentiation capability. LT-HSCs first differentiate into short-term hematopoietic stem cells (ST-HSCs) and then multipotent progenitor cells (MPPs). MPP cells differentiate into either common lymphoid progenitor (CLP) or common myeloid progenitor (CMP) cells. CMPs then further branch out into granulocyte-macrophage progenitors (GMPs) and megakaryocyte-erythrocyte progenitors (MEPs). The GMP cells are then terminally differentiated into granulocytes (GR), while MEP cells are further differentiated into megakaryocyte progenitors (MKP) and then terminally differentiated into megakaryocytes (MK).

**Figure 6:**
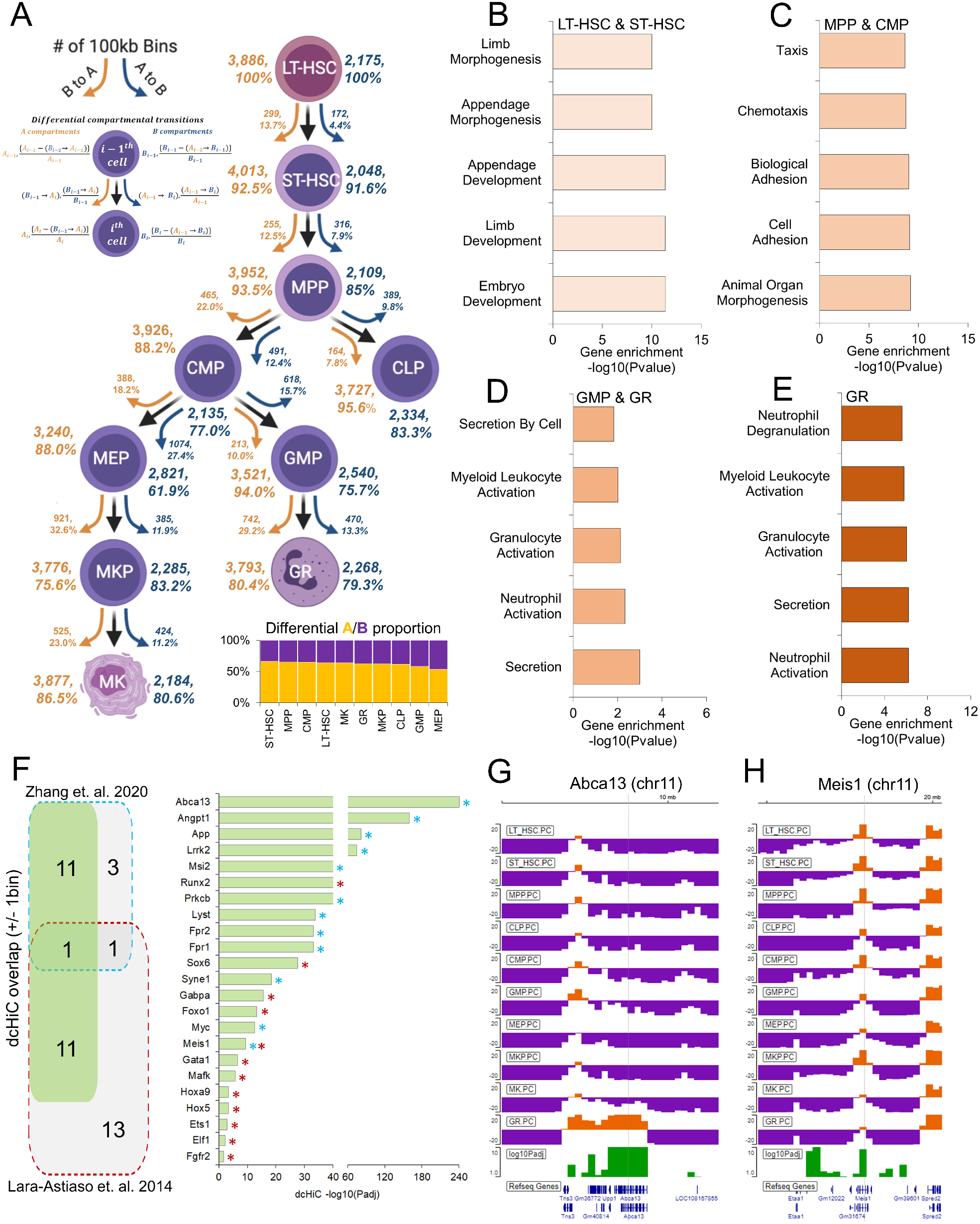
Ten-way multivariate differential compartment analysis of mouse hematopoiesis. **(A)** Summary of overall compartment decomposition and significant compartment changes observed across the 10 cell types. The orange and blue arrows represent A to B and B to A compartment flips, respectively. The numbers next to the arrows represent the total number of flipping compartments, and the numbers within the parentheses next to arrows show the significantly different flipping compartments. The bottom-right plot shows the proportion of A and B bins among *dcHiC* differential compartments for each cell type. Figure adopted from Zhang et al. [28]. **(B-E)** The functional enrichment of genes overlapping with differential compartments from 10-way comparison that have the strongest A compartment scores in either **(B)** LT-HSC or ST-HSC, **(C)** MPP or CMP, **(D)** GMP or GR, or **(E)** GR alone. **(F)** Differential compartments identified by *dcHiC* overlap a set of critical genes previously known to play a role in mouse hematopoiesis. **(G)** An IGV browser snapshot of the *Abca13* gene and its overlapping differential compartment across the ten cell types. The *Abca13* gene is exclusively found to be a part of the A compartment in GR but in the B compartment in all other cell types. (**H)** An IGV browser view surrounding the *Meis1* genic region. This region overlaps with the A compartment for all cell types but with varying magnitudes of strength. *dcHiC* captured this region as a differential compartment.

Using the Hi-C data from this system, we carried out multivariate differential analysis using *dcHiC* at 100 kb resolution. We detected a total of 6,061 (60.61 Mb of the genome) differential compartment bins across the ten cell types encompassing many of the genomic regions previously shown to undergo hematopoiesis-related dynamic changes [28]. **Figure 6A** shows an overall summary of the significant compartment changes identified by *dcHiC* across these cell types. We observed that the number of A to B transitions continued to increase from the LT-HSC stage to the MEP and GMP progenitor stages. The differentiation of CMP into MEP and GMP cells represents two of the most frequent A to B transitions (∼27.4% and ∼15.7% A → B transition, respectively) within the hematopoietic hierarchy, likely reflecting the need for suppression of certain transcriptional profiles for commitment into each branch. This is consistent with the largest proportion of differential B compartments in MEP (∼46.5%) and GMP (∼42%%) compared to all other cell types. With respect to the top of the hematopoietic tree (i.e., LT-HSC), early progenitors such as MPP have 571 100 kb bins with a significant compartment flip (either A to B or B to A), whereas the differentiated cells such as MK and GR had 949 and 1,212 such bins, respectively. This confirms the gradual divergence of chromatin compartmentalization from hematopoietic stem cells as cells progress further into differentiation.

Next, similar to the ESC-NPC-CN transition, we also carried out functional enrichment analysis of differential regions with the highest A compartment score in each group and specific cell type. **Figure 6B-E** shows these enrichments for four different stages of hematopoiesis (pre-bifurcation stage: LT-HSC, ST-HSC, progenitor stage: MPP, CMP, granulocyte branch: GMP, GR and the terminally differentiated granulocytes or GR) with respect to the rest and for a specific cell type within each of these stages highlighting biologically relevant processes in each case. For example, morphogenesis- and development-related biological processes were enriched in the overall prebifurcation stage (set of genes with the highest A compartment score in either LT-HSC or ST-HSC) (**Figure 6B**), and progenitor stage cells were enriched in morphogenesis-, adhesion- and migration-related terms (**Figure 6C**). The granulocyte branch (GMP and GR) as well as the terminally differentiated granulocytes (GR) showed significant enrichment related to the activation and regulation of neutrophils and granulocytes (**Figure 6D-E**). For the megakaryocyte branch (MEP, MKP, MK), however, we did not observe any statistically significant GO term biological process enrichment.

#### Example genes from mouse hematopoiesis differential compartments

After investigating the significance of the differential compartments from a high level, we examined the genes overlapping with the differential compartments involved in hematopoietic lineage differentiation and chromatin dynamics [54]. **Figure 6F** shows a set of important genes overlapping with differential compartments from our multivariate analysis. Zhang et al. showed that increased gene-body associating domain (GAD) scores are linked to active transcription and indicate cell-type specific features. We identified 12 out of 16 such differential GAD genes between ST-HSC and GR as part of *dcHiC* differential compartments identified across the system (FDR < 0.1; **Figure 6F**, marked by cyan stars). In addition, a previous analysis by Lara-Astiaso et al. [54] also reported a set of critical genes for hematopoietic lineage differentiation. We identified 12 of these 26 genes within differential compartments (FDR < 0.1; **Figure 6F**, marked by red stars), supporting *dcHiC’s* ability to detect changes in regions harboring genes that are dynamically regulated during hematopoiesis. Among these genes, one example is the transmembrane transporter gene *Abca13*, which was the exclusive differential A compartment within GR but in the B compartment for all other cell types **(Figure 6G)**. Other notable examples include *Meis1*, a transcription factor required to maintain hematopoiesis under stress and over the long term [55]. Notably, this particular example was a significant change solely within the A compartment **(Figure 6H)**. Apart from *Meis1*, dcHiC also detected differences for other transcription factors, such as *Runx2* and *Sox6*, that are essential for progenitor cell differentiation (**Figure 6F)** [56, 57]. We also identified *Myc*, known for its role in balancing haematopoietic stem cell self-renewal and differentiation [58] adjacent to a significant change within the A compartment that encompasses the *Pvt1* gene. The long noncoding RNA *Pvt1* harbors intronic enhancers that interact with *Myc* and promote *Myc* expression during tumorigenesis [59]. Overall, this complex system demonstrates the utility of *dcHiC’s* multivariate compartment analysis, which discovers important changes in compartmentalization without requiring a large number of pairwise comparisons.

### Multiway differential compartment analysis across human-derived cell lines

Measuring the extent to which genetic variation across individuals influences chromatin features, including 3D organization, has significant implications in our understanding of human disease. Previous studies have revealed that the presence of variations such as quantitative trait loci (QTLs) can affect histone modifications, transcription factor binding, and enhancer activity across populations [60, 61]. More recent work by Gorkin et al. [62] studied variation in chromatin conformation across individuals from different human populations. Using dilution Hi-C, they profiled lymphoblastoid cell lines (LCLs) derived from 13 Yoruban individuals, one Puerto Rican trio, one Han Chinese trio, and one European LCL (GM12878). They measured significant differences in 3D genome organization across individuals using different metrics, including the Directionality Index (DI), Insulation Score (INS), Frequently Interacting REgions (FIREs), and compartment scores [62]. The study also carried out differential analysis of compartments across individuals and provided both compartment scores and “variable regions” at 40 kb resolution (except for chromosomes 1, 9, 14, 19 and X). To minimize technical variation and ensure a fair comparison, we started directly from the 40 Kb compartment scores reported by Gorkin et al. and ran *dcHiC* on these values (starting from quantile normalization). *dcHiC* allows direct utilization of precomputed compartment scores, such as in this case, when available.

The Venn diagram (**Figure 7A**) of differential compartments from *dcHiC* and Gorkin et al. using the same set of 40 kb genomic bins shows a large overlap between the methods. A large fraction of *dcHiC* calls (7524 out of 7,876 or ∼96%) were also reported by the original paper. However, Gorkin et al. reported an additional 765 Mb of the human genome as variable compartment regions (**Additional_file_4.xlsx** from the original publication filtered for phenotype=PC1 and discover_set=20 LCLs), which amounts to ∼11K more bins at 40Kb resolution. To further study the overlap and differences between the two approaches, we plotted two statistical significance score distributions (-log10 of the adjusted p-value calculated by Gorkin et al.) for regions that the Gorkin study reported as differential, one with regions overlapping with *dcHiC* calls and the other with nonoverlapping regions (**Figure 7B**). Variable compartments from the previous study that were not deemed significant by *dcHiC* have substantially lower statistical significance, as computed by the original paper suggesting *dcHiC* calls are enriched for stronger differences. Next, we compared the full set of differential compartments called by both methods and their fraction covering each individual chromosome **(Figure 7C)**. The figure shows that Gorkin et al. calls cover a larger fraction of smaller chromosomes, with more than half the entire length reported as a significant variable compartment for some chromosomes (e.g., chr18). *dcHiC*, on the other hand, has a more uniform representation of differential compartments across chromosomes, with differential fractions ranging between 10% and 20% for most chromosomes. Finally, we compared the top 5000 differential compartment bins ranked by their significance scores from each approach. **Figure 7D** shows that ∼61% of these top 5000 differential bins are identical, suggesting substantial differences in each approach’s ranking with respect to statistical significance (Spearman rank correlation of 0.55). Although the ranking is substantially different between the methods, the overlapping fraction of the top 5000 differential compartments for *dcHiC* had more significant differences (**Figure 7E**). Using other variable chromatin organization metrics from the Gorkin paper, we observed that *dcHiC* calls were more enriched in FIRE-QTLs **(Figure 7F)** as well as DI-QTLs **(Figure 7G)**. Preferential enrichment of such signals suggests a better concordance of *dcHiC* identified compartmental differences and chromatin organization variability at other levels across individuals.

**Figure 7:**
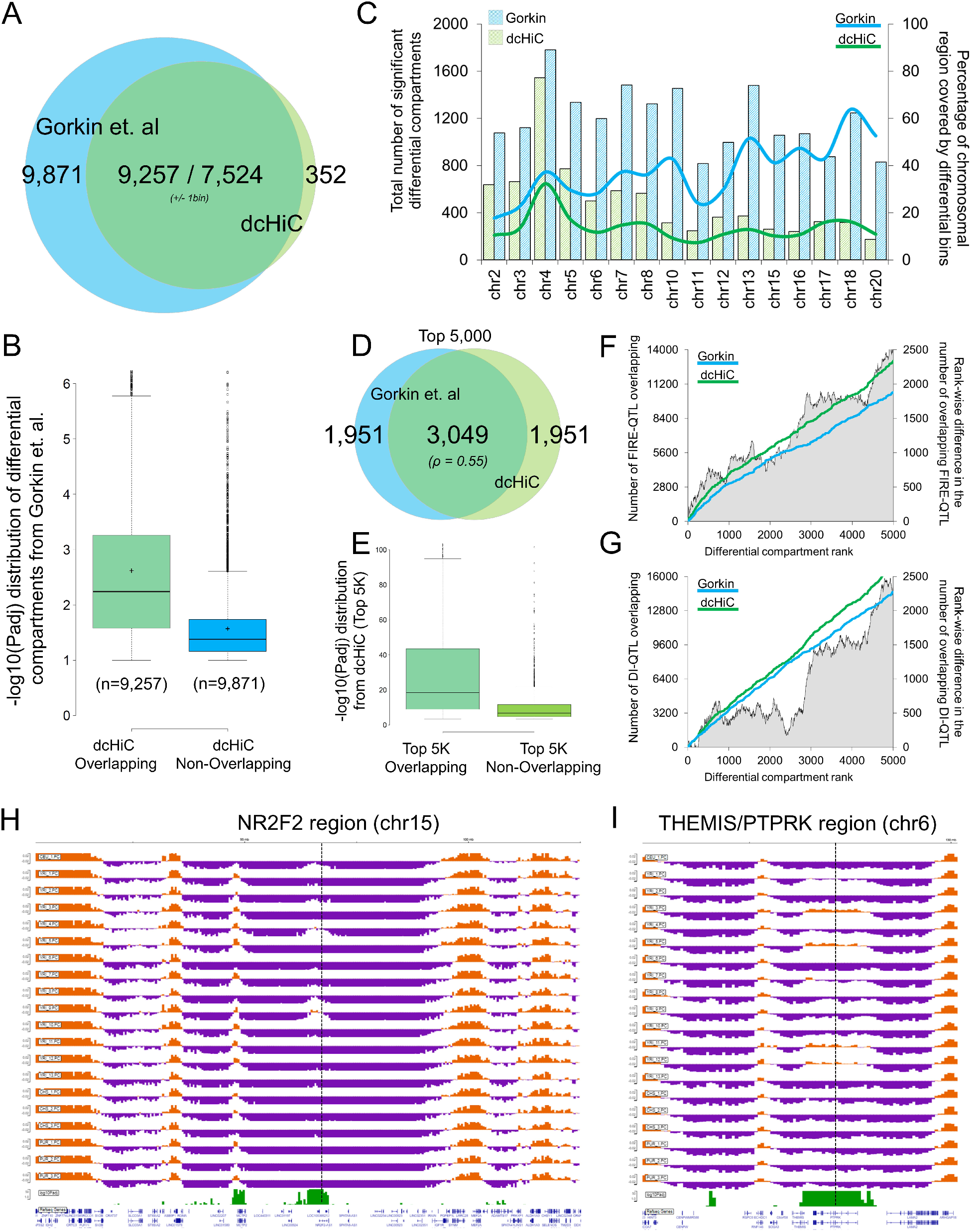
Twenty-way multivariate differential compartment analysis of human lymphoblastoid cell lines (LCLs). **(A)** A Venn diagram of the overlap of differential compartments called by *dcHiC* and the *variable compartment regions* from a previous study (Gorkin et al., 2019). **(B)** The distribution of -log10(p-adj) values of *dcHiC*-overlapping and non-overlapping variable regions calculated by the previous study. **(C)** The total number of chromosome-wise differential compartments and the fraction of each chromosome (except those filtered by Gorkin et al. 2019) covered by such calls for *dcHiC* and the previous study. **(D-E)** Venn diagrams of overlapping compartments of the top 5000 differential regions from both approaches and the –log10(p-adj) value distribution of the overlapping and nonoverlapping sets from *dcHiC*. **(F-G)** The cumulative number of FIRE-QTLs and DI-QTLs overlapping the top 5000 differential compartment calls by *dcHiC* and Gorkin et al. **(H-I)** Two differential compartments overlapping genic regions of *NR2F2* and *THEMIS/PTPRK*. Both the genes and especially the *NR2F2* region were shown to be variable regions across the population through FISH experiments in a previous study (Gorkin et al., 2019).

#### Example genes from differential compartments among human-derived LCLs

**Figures 7H-I** show two examples of a variable region (overlapping with *NR2F2* and *THEMIS/PTPRK* genes) identified by *dcHiC*. The *NR2F2* region was investigated using FISH by Gorkin et al., which confirmed individual-specific changes in 3D chromatin conformation. Two of the individuals from the cohort (YRI-4 and YRI-8) showed enriched interaction between the *NR2F2* FISH and another placed upstream compared to YRI-3 and YRI-5. The variability of 3D genome organization among individuals is also apparent from compartment scores for this region. The *NR2F2* locus across the cohort was found to be a part of a strong B-compartment for all Yoruban individuals except for YRI-4, YRI-8 and YRI-9 **(Figure 7H). Figure 7I** shows another example of such a variable region with coordinated changes in epigenetic marks across individuals with support from differential compartments documented in a previous paper. Gorkin et al. identified variations in different epigenetic marks, such as H3K4me1 and H3K27ac, binding of CTCF and, most importantly, gene expression patterns within this region across different individuals (YRI-2 and 13 vs 11 and 12). The PC score track in **Figure 7I** also supports the previous findings, as some of the individuals from the YRI population, especially YRI-3, YRI-5, YRI-11, and YRI-12, showed a clear flip from the B to A compartment, and both our approach and Gorkin et al. labeled this region as a differential compartment. Taken together, dcHiC identified fewer differential compartment bins with enrichment towards capturing regions with higher variability in different levels of chromatin organization and those with additional evidence for differences among individuals.

## DISCUSSION

This paper presents a new application, *dcHiC*, to compare compartmentalization across Hi-C datasets. *dcHiC* employs principal component analysis followed by quantile normalization of the compartment scores and a multivariate distance measure to systematically identify significant compartmentalization changes among multiple contact maps. By facilitating comparative analysis across multiple integrated datasets, it helps identify biologically relevant differential compartments with statistical confidence scores. Along with conventional pairwise differential analysis, *dcHiC* allows a single multivariate differential comparison of Hi-C datasets, utilizing replicates when available, and provides an efficient approach to analyze multiple Hi-C maps without the need for generating many different combinations.

We applied *dcHiC* to various biological scenarios, ranging from neuronal and hematopoietic stem cell differentiation in mice to Hi-C data from different human populations. Our results confirmed that *dcHiC* detects known compartmental changes among cell types, including those previously validated to play a role in neuronal and hematopoietic differentiation. When comparing *dcHiC* to existing approaches, we showed that it identifies regions with higher differences in replication timing, Lamin B1 signals, and differentially expressed genes, suggesting better prioritization of relevant biological regions. Even though differences in compartmentalization between ESCs and NPCs are generally aligned with changes in the lamin B1 association, a recent work highlighted the importance of nucleolus association in revealing layers of compartmentalization with distinct repressive chromatin states [63]. Our initial analysis showed that over 10% of all significant compartment differences we found between ESC and NPC belong to nucleolus-associated domains (NADs) that were deemed exclusive to either ESC or NPC [63], providing an explanation for a subset of differences in compartmentalization during differentiation. Expanding on a three-way (n=3) mouse neuronal differentiation model, we showed that *dcHiC* continues to systematically identify critical biological marker genes and can recover cell-specific functions from differential compartment analysis alone. Across *dcHiC’s* differential compartments, we observed significant and relevant enrichment of biological processes such as neuron differentiation in NPC and CN cells. More broadly, *dcHiC’s* differential compartments also compellingly aligned with changes in Lamin B1, gene expression, and histone modification data. Taken together, these results demonstrate *dcHiC’s* ability to find regions with the most biologically variable compartmentalization across the genome.

The hierarchical mouse hematopoietic stem cell differentiation model, consisting of ten different cell types with Hi-C data, provided a unique opportunity to demonstrate the utility of *dcHiC*. A ten-way multivariate differential comparison of the hematopoietic system revealed previously known lineage-specific critical genes encompassing the differential compartments. Notably, we identified vital transcription factors, such as *Sox6, Runx2, Meis1, Foxo1*, and many other critical genes, such as *Abca13*, by solely analyzing the differential calls. Our functional enrichment analysis of gene sets overlapping with the lineage-specific differential compartments reported from the apex to the bottom of the hematopoietic model tree reconfirmed that genome compartments play a contributory role in determining the accessibility of genes in specific cell types. Measuring the extent of compartment variability across twenty-cell human types also highlighted our method’s novel utility and strength. Most *dcHiC* calls overlapped with a subset of variable compartments reported by the previous study, but *dcHiC* calls were enriched for higher variability. Similarly, the regions encompassing Frequently Interacting Region (FIRE)-QTLs and Directionality Index (DI)-QTLs defined by the previous study were more enriched in the top differential compartment calls of *dcHiC* than in the top calls defined in the previous study. The analysis also demonstrated an important feature of *dcHiC*: the ability to directly utilize previously computed compartment scores to run differential compartmentalization analysis.

The framework we developed here provides a systematic way to identify differential compartments and visualize these differences in different scenarios, including multiway, hierarchical and time-series settings. Although we focused on human and mouse Hi-C data in this work, our method is readily applicable to Hi-C data or its variants (e.g., Micro-C [64]) derived from any organism with compartmental genome organization. With hundreds of publicly available Hi-C datasets in the 4D Nucleome Data Portal and others published every day, *dcHiC* will play an essential role in the comparative analysis of high-level genome organization. As single-cell Hi-C data start providing better resolution, *dcHiC* and methods derived from it will be critical to enable compartment comparison across thousands of cells.

## METHODS

### Data processing, result generation, and visualization

#### Hi-C data

All the Hi-C data, except for Gorkin et al. 2019, were mapped to the mm10 reference genome and processed using the HiCpro (v2.7.9) pipeline [65]. The raw Hi-C interaction maps retrieved after HiC-Pro processing are used for downstream compartment score calculation by *dcHiC*. In the section analyzing data from the Gorkin et al. study, we used the provided compartment scores (40 kb resolution) across all samples mapped to the hg19 reference genome [62]. Statistically significant interactions were called using FitHiC2 [44] with default parameters and an FDR threshold of 0.05 for each replicate and/or each sample.

#### RNA-seq data

The RNA-seq data from Bonev et al. 2017 [38] study concerning mouse neural development were processed using our in-house and open-source RNA-seq processing pipeline (https://github.com/ay-lab/LJI_RNA_SEQ_PIPELINE_V2.git), which utilizes STAR [66]. The differential gene expression analysis between mouse ESC and NPC cell lines (two replicates each) was performed using the DESeq2 method [39] with all the default parameter settings.

#### ChIP-seq data

For ChIP-seq peak calling (H3K27ac, H3K4me3 and H3K4me1 histone marks), we first mapped the respective fastq files to the mm10 genome using bowtie2 [67] and generated the corresponding bam files (MAPQ > 20). The aligned files were then used as input to the MACS2 program [68] to call peaks (p-value < 1e-5) against their respective input controls. The continuous ChIP-seq peaks were then merged, and the unique set was mapped to the 100 kb differential compartments to calculate the average number of peaks. The enrichment of signal difference was calculated by first quantifying the absolute difference in signal (number of ChIP-seq peaks and gene expression TPM values) within ESC to NPC differential and nondifferential compartments. The enrichment of absolute signal difference between the differential and non-differential compartments between ESC and NPC was then compared by un-paired T-test.

#### Time-series analysis

Time-series clustering was generated using the TCseq package [46]. For gene-term enrichment analysis, the differential compartments are scanned against the gene coordinates of the respective genome defined by the user using the ‘bedtools map’ function [69]. The unique overlapping set of genes was then extracted and used for GO biological function enrichment analysis using the ToppGene suite API function or directly from their webserver [47].

#### IGV Browser visualization

*dcHiC* generates a JavaScript-based stand-alone dynamic IGV-HTML page to visualize the compartments and differential compartment calls, with an option to add additional tracks.

### Computation and quantile normalization of compartment scores for comparison

To perform principal component analysis (PCA) on Hi-C maps, *dcHiC* utilizes the singular value decomposition (SVD) implementation of the bigstatsr R package [32]. The input to SVD is *K* different distance-normalized chromosome-wise correlation matrices *(X*_1_, *X*_2_, *X*_3_…*X*_*K*_) for each Hi-C data. For each such matrix, *dcHiC* finds the decomposition:

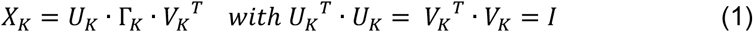

The matrices *U*_*K*_ and *V*_*K*_ store the left and right singular vectors of the matrix *X*_*K*_. The singular values of *X*_*K*_ are stored in the diagonal matrix Γ_*K*_. The principal components for each matrix are then obtained as:

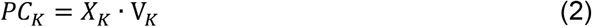

The eigen-decomposition of the *K*^*th*^ correlation matrix provides the eigenvectors, and the sign of the first eigenvector or principal component (*PC*1_*K*_) typically represents the genomic compartments A and B for the *K*^*th*^chromosome. If *PC*1_*K*_ corresponds to chromosome arms or other broad patterns in the Hi-C matrix, the second principal component (*PC*2_*K*_) may represent A and B compartments. The A/B compartment labels are assigned to the positive/negative stretches of the selected *PC*_*K*_ depending on the implementation of eigen-decomposition. It may be necessary to reorient these assignments and select the correct *PC*_*K*_ using GC content or gene density. Thus, before the quantile normalization step, *dcHiC* performs an intermediate correlation analysis of the first two principal component scores (user-defined) of each chromosome per sample against the GC content and gene density of that chromosome. The principal component that obtained the highest sum of GC content and gene density correlation was considered the compartment score, and the A/B compartments of the selected principal components were assigned based on the GC content correlation (A compartment and positive values representing higher GC content). These generate a set of compartment score vectors representing each sample (*M* samples) for a given chromosome *(C*_1_, *C*_2_, *C*_3_ … *C*_*M*_). Once the properly labeled compartment scores are obtained, *dcHiC* performs quantile normalization (QN) using the limma package [70] on the set *(C*_1_, *C*_2_, *C*_3_ … *C*_*M*_) per chromosome to even out the scaling across the group for downstream analysis.

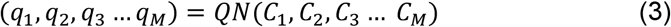

In the case of samples with replicates, *dcHiC* performs the above steps by including each replicate from each sample (i.e., quantile normalizes all replicates together). *dcHiC* then calculates the average quantile normalized values of each genomic bin across all replicates of a given sample to represent sample-wise compartment scores.

### Differential compartment identification

Mahalanobis distance (MD) is a multivariate statistical measure of the extent to which the multivariate data points are marked as outliers based on a Chi-square distribution [71]. The Mahalanobis distance of a point *i* from a multidimensional distribution defined by set *s (sample)* and its center *µ* is defined as:

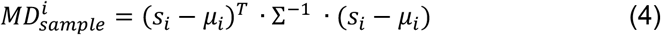

where 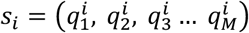 is the set of quantile normalized compartment score distributions and 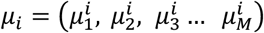 is the set of weighted centers for each point *i* from set *s*. The inverse of the covariance matrix of set *s* is represented as Σ^−1^. The weighted centers *µ*_*i*_ are calculated as:

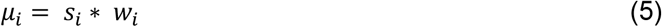

where 0 ≤ *w*_*i*_≤ *1* is the cumulative normal distribution probability associated with the maximum z-score among the z-scores of all samples for *i*:

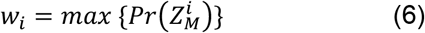

where 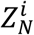, the z-score for point *i* for sample *N*, is computed as:

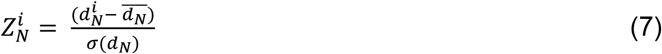

Here, 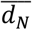 and *σ(d*_*N*_) represent the average distance and standard deviation within sample *N* among all 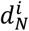 values that are computed as:

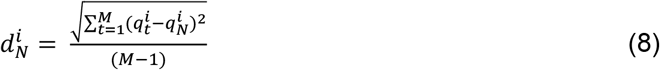

Essentially, the approach provides more weight to the points that are distant from others among the samples (further from the diagonal) than to points that are closer together in the multidimensional space (close to the diagonal). Equation (4) is the standard MD formulation, which we modify using the weighted centers as computed through Equations (6) to (8).

To increase the sensitivity of our difference detection, we implemented an outlier removal step that eliminates genomic bins (or points) with high MD (as computed above) at the initial pass (1^st^ pass). We use a predefined upper-tail critical value of the chi-square distribution with *df* degrees of freedom as our threshold for outlier removal (the default value we used is 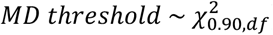). We then recompute the covariance matrix Σ^−1^ after removal of these outliers and calculate the MD (through Equation (4)) one more time for each point (2^nd^ pass). The significance of the corresponding 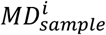 (2^nd^ pass) is calculated from the critical chi-square distribution table as 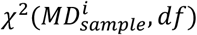 using the *pchisq* function of the R programming language followed by multiple testing correction to retrieve adjusted p-values.

In the case of samples *(s)* with replicates, *(r*) *dcHiC* calculates an additional covariate *MD*_*replicate*_ and applies Independent Hypothesis Weighting (IHW) to adjust the *p-values*.

The covariate is calculated as follows:

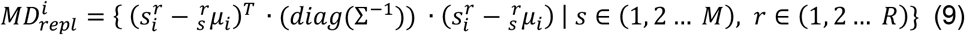

where *R* is the total number of all replicates combined across all samples, 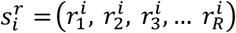 is the set of quantile normalized compartment score distributions of all replicates from samples *s* ∈ (1, 2 … *M*) and 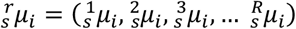 is the set of weighted centers for each point *i* from *R* replicates. *diag* is an operation that masks all nondiagonal entries (sets to zero) of the covariance matrix.

The weighted centers 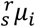 are calculated as:

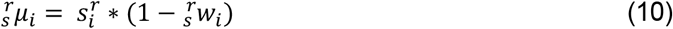

where 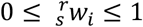 is calculated as:

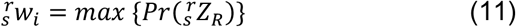

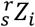 for replicate *r* of sample *s* is computed as:

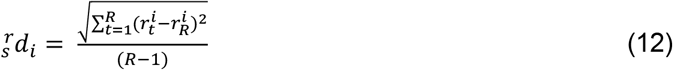

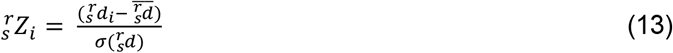

Here, the variables are defined as similar to Equations (7) and (8), and *R* is used to represent the number of replicates of the same sample (i.e., distances across replicates of different samples are not taken into account).

This approach provides more weight to the features that are closer to each other within replicates of a sample (close to the diagonal) and as opposed to the calculation across different samples (Equation (5)), where higher weights were given to the points with samples distant from each other (far from the diagonal). The significance of the corresponding *MD* for each point is calculated using the chi-square distribution asc mentioned above. *dcHiC* applies the IHW approach to adjust the p-values using FDR correction obtained from *MD*_*sample*_ using the *MD*_*replicate*_ replicate variation measure as a covariate.

### Differential interaction identification

Using the same Mahalanobis distance (MD) measure, *dcHiC* enables the user to find differential interactions across samples that are either linking two differential compartments together or a differential compartment with other parts of the same chromosome. The goal of this feature is to provide more information on the chromatin organization changes related to or correlated with compartmental differences. For this analysis, we used FitHiC2 to call significant interactions (FDR 5%) for each sample or replicate (when available), but users are free to provide their own set of interaction or loop calls from any other tool. Using these calls, *dcHiC* first finds the interaction subset that overlaps with differential compartments (on either end or both) using the bedtools *‘pairtobed’* function. *dcHiC* utilizes the 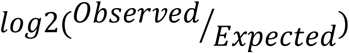 values of a chromatin interaction *i* to perform differential interaction calling as follows:

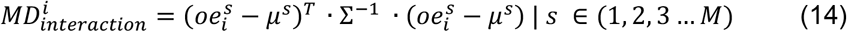

where 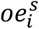 represents 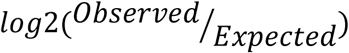 values for chromatin interactions of locus pair *i*for sample *s* and *µ*^*s*^ represents the vector of centers of distance normalized interactions from sample *s*. Here, Σ^−1^ represents the inverse of the covariance matrix of interactions among the samples. The approach provides more weight to the interactions that are distant from the expected interaction strength among the samples than to the interactions that are closer to the expected range in the multidimensional space. The significance of the corresponding 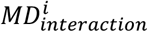 is calculated from the critical chi-square distribution table as 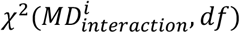 using the *pchisq* function embedded within the R programming environment followed by FDR correction to retrieve adjusted p-values.

## Availability of data and materials

A Python/R implementation of *dcHiC* is freely available at https://github.com/ay-lab/dcHiC. This application is compatible with Hi-C data in HiC-Pro, .*hic*, and .cool formats. The data used in this study are available at the following GEO accession numbers: GSE96107 (mESC-NPC-CN), GSE152918 (mouse hematopoiesis), and GSE128678 (human LCLs). These are also available in **Supplemental Table 1** (see Supplementary Information). All reported compartments for all cell lines, multivariate differential scores, RNA-seq, and ChIP-seq data used in this manuscript can be viewed interactively at ay-lab.github.io/dcHiC. These standalone HTML files employ *dcHiC’s* visualization utility through the IGV browser.

