## Supplementary Information for "dcHiC: differential compartment analysis of Hi-C datasets"

**Supplemental Information**

**Table S1: Publicly available datasets used in this paper**

| **Accession** | **Data Type** | **Cell Lines** | **Factor(s)** | **Reference** |
| --- | --- | --- | --- | --- |
| GSE96107 | Hi-C | mESC, NPC, CN | None | Bonev et. al (2017) |
| GSE96107 | RNA-Seq | mESC, NPC, CN | None | Bonev et. al (2017) |
| GSE96107 | ChIP | mESC, NPC, CN | H3K27ac, H3K4me1, H3K4me3 | Bonev et. al (2017) |
| GSE152918 | Hi-C | LT-HSC, ST-HSC, MPP, CMP, GMP, GR, MEP, MK | None | Zhang et. al (2020) |
| GSE128678 | Compartment scores | Human population cohort | None | Gorkin et. al (2019) |

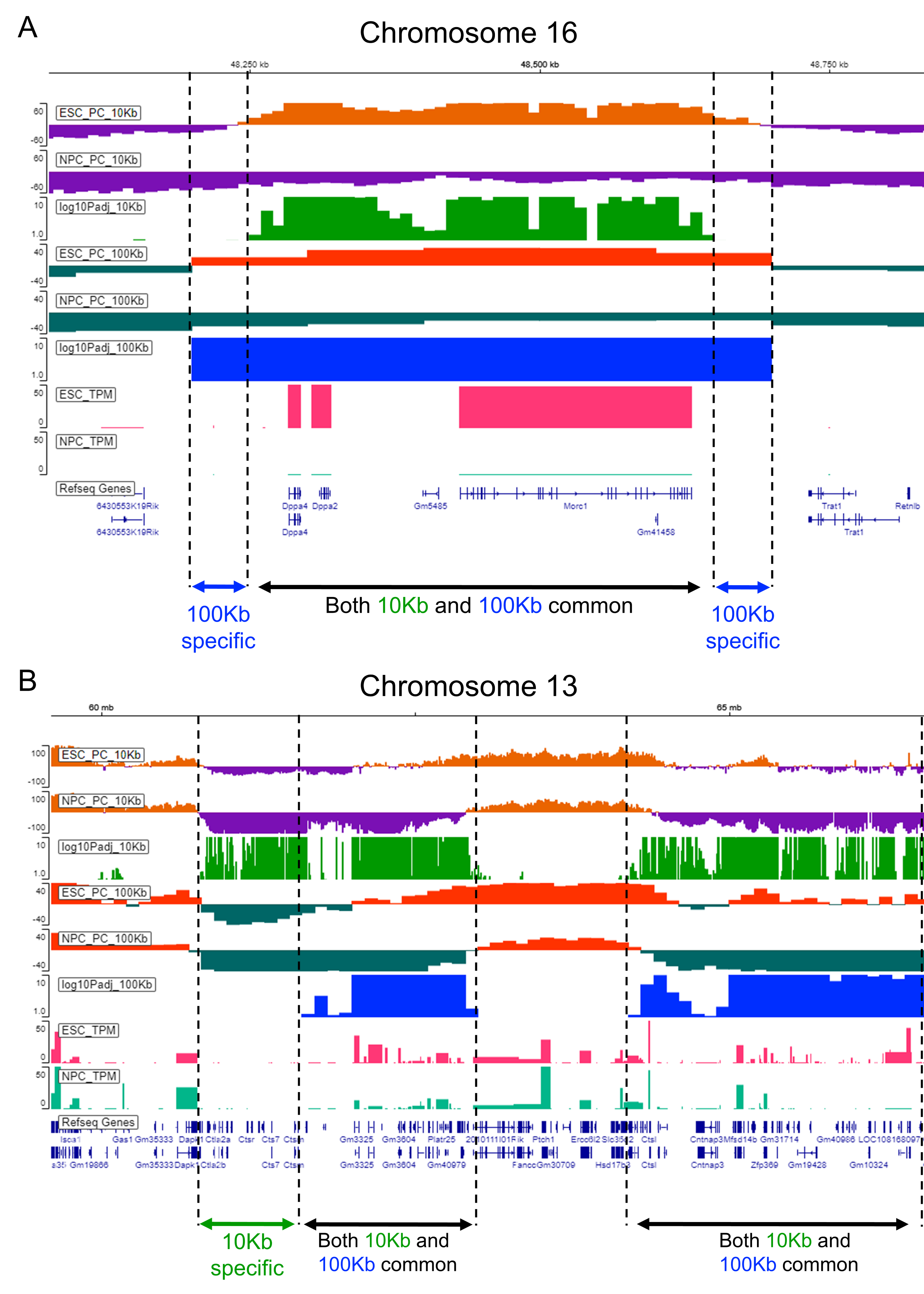

**Supplementary Figure 1: Comparison of 100Kb and 10Kb dcHiC analysis of ESC – NPC compartments. (A)** Genomic region containing Dppa2/4 genes with 10kb vs 100kb differences at the boundaries of common differential compartments. **(B)** A chromosome 13 region with differential compartments that are identified only at the 10Kb resolution *dcHiC* analysis.
